# Embryonic Exposure to Valproic Acid Disrupts Social Predispositions in Newly-Hatched Chicks

**DOI:** 10.1101/150391

**Authors:** Paola Sgadò, Orsola Rosa-Salva, Elisabetta Versace, Giorgio Vallortigara

**Author notes:** These authors contributed equally to this work (Orsola Rosa-Salva, Elisabetta Versace). Address for correspondence: Paola Sgadò, PhD, University of Trento - Center for Mind/Brain Sciences Animal Cognition and Neuroscience Laboratory Piazza della Manifattura 1, 38068 Rovereto (TN), Italy, Phone: ©39 0464 808691.

## Abstract

Biological predispositions to attend to visual cues, such as those associated with face-like stimuli or with biological motion, guide social behavior from the first moments of life and have been documented in human neonates, infant monkeys and newly-hatched domestic chicks. In human neonates at high familial risk of Autism Spectrum Disorder (ASD), a lack of such predispositions has been recently reported. Prompted by these observations, we modeled ASD behavioral deficit in newborn chicks, using embryonic exposure to valproic acid (VPA), the histone deacetylases (HDACs) inhibitor that in humans is associated with an increased risk for developing ASD. We assessed spontaneous predispositions in newly-hatched, visually-naïve chicks, by comparing responses to a stuffed hen *vs.* a scrambled version of it. We found that social predispositions were abolished in VPA-treated chicks. In contrast, experience-dependent learning mechanisms associated with filial imprinting were not affected. Our results indicate a specific effect of VPA on the development of biologically-predisposed social orienting mechanisms, opening new perspectives to investigate the molecular and neurobiological mechanisms involved in early ASD symptoms.

## Introduction

Autism Spectrum Disorder (ASD) comprises a genetically heterogeneous group of neurodevelopmental disabilities characterized by a wide range of impairments in social behaviors. Given their genetic heterogeneity and the complex behavioral traits associated with ASDs diagnosis, animal models are essential for the study of the mechanistic bases of these disorders and for development of potential therapies. Despite several studies addressing the importance of early diagnosis and intervention in ASD [1], to date symptoms recognition is achieved after 2 years of age, limiting the possibility of early treatments. Delineating the earliest expression of ASD would not only increase the opportunities for intervention but also advance our understanding of the underlying biology, thus generating new therapeutic options. Hence, establishing animal models to investigate the early development and mechanistic bases of ASD could have a crucial impact on the development of therapies for ASD.

One of the aspects that limit neurodevelopmental studies in existing animal models of ASD is the availability of early social behavioral tests, reliably recapitulating the social impairment shown in the patients. Biological predispositions to attend to social stimuli, without any previous experience (social predispositions thereafter), are the earliest expression of social behavior. Social predispositions have been described in humans [2], non-human primates [3] and domestic chicks [4], as spontaneous, hard-wired, mechanisms that drive visual attention to features associated with social partners [5]. Typical newborn babies show, for instance, preference for faces and face-like configurations [5] exactly as newly-hatched chicks [6,7] and naïve infant monkeys do [8]. These inter-species similarities in response to social cues (or more generally to cues of “animacy”) extend to biological motion [9,10], self-propulsion [11,12], and speed changes [13] (though in some cases species differences have been also reported [14]). Most important, some of the subpallial areas linked to social predispositions have started to be identified [15-17].

Social predispositions have also been associated with behavioral deficits in ASD. A recent prospective study analyzed these early-emerging mechanisms in newborn babies (four to ten days old) with a high familial risk of ASD [18]. Measuring visual attention towards face-like stimuli and biological motion cues, an impairment in the orienting mechanisms towards these social stimuli was observed in neonates at high-risk for ASD, compared to the typical population. This discovery moves the potential for early ASD assessments to the first moments of life, thus increasing the interest for models reproducing symptoms that can be assessed soon after birth.

Clinical studies have shown that prenatal exposure to the histone deacetylases (HDACs) inhibitor valproic acid (VPA) is associated with neural tube malformations, reduced cognitive function and an increased risk for developing ASD [19]. VPA directly inhibits HDACs [20], interfering with normal deacetylation of chromatin and causing activation of aberrant gene transcription during development [21]. Given the strong association of VPA treatment with development of social behavioral deficits in humans, animal studies using prenatal exposure to VPA have been conducted, to model the core signs of ASD and to identify the molecular pathways linked to ASD social deficits [22,23]. Despite several investigations devoted to the study of histone acetylation and the effect of HDAC inhibitors on memory and cognition [24,25], the detrimental effect of these compounds on brain development and early social behaviors, and their role in the etiology of ASD is still unclear. Previous studies conducted in chicks showed that VPA can alter aggregative behavior and decrease vocalizations [26]. To investigate the contribution of social predispositions to atypical social behavior related to ASD in an animal model that allows for controlled experimental conditions, we delivered VPA *in ovo*, in the last week of embryogenesis, and compared the performance of VPA- and vehicle-injected chicks on social predispositions to approach a social stimulus (a stuffed hen), and on affiliative responses mediated by the learning mechanism of filial imprinting.

## Results

Previous studies demonstrated that dark-hatched chicks prefer to approach a stuffed hen over an artificial object, or even over a scrambled version of the same stimulus [27]. We performed the same test in visually-naïve VPA- and vehicle-injected domestic chicks and assessed their predisposed preference to approach a naturalistic stimulus consisting of a stuffed hen over a “non-social” stimulus, in which features of the hen were dismantled and attached on the sides of a box in scrambled order [27] (see Methods for details, see Figure 1A). To evaluate social predispositions we calculated the preference score for the predisposed stimulus as the proportion of running wheel revolutions toward the stuffed hen (see Methods for details). Results showed an effect of treatment on the chicks’ preference for the predisposed stimulus. While in the control group the preference for approaching the stuffed hen was similar to what previously observed [16,28] (Figure 2A), VPA treatment significantly reduced the preference for the stuffed hen compared to controls (Figure 2A). In line with this observation, the average preference score for the predisposed stimulus was significantly different from chance level only for the control group and not for VPA-treated chicks (Figure 2A). Thus, VPA treatment significantly reduces the predisposed preference for the stuffed hen to chance level. On the contrary, we did not observe significant differences between treatments in motor activity, measured as the overall number of rotations in the running wheel (Figure 2B). We detected however a difference between the two sex groups: female were significantly more active than males, irrespective of treatment (Figure 2C).

**Figure 1.**
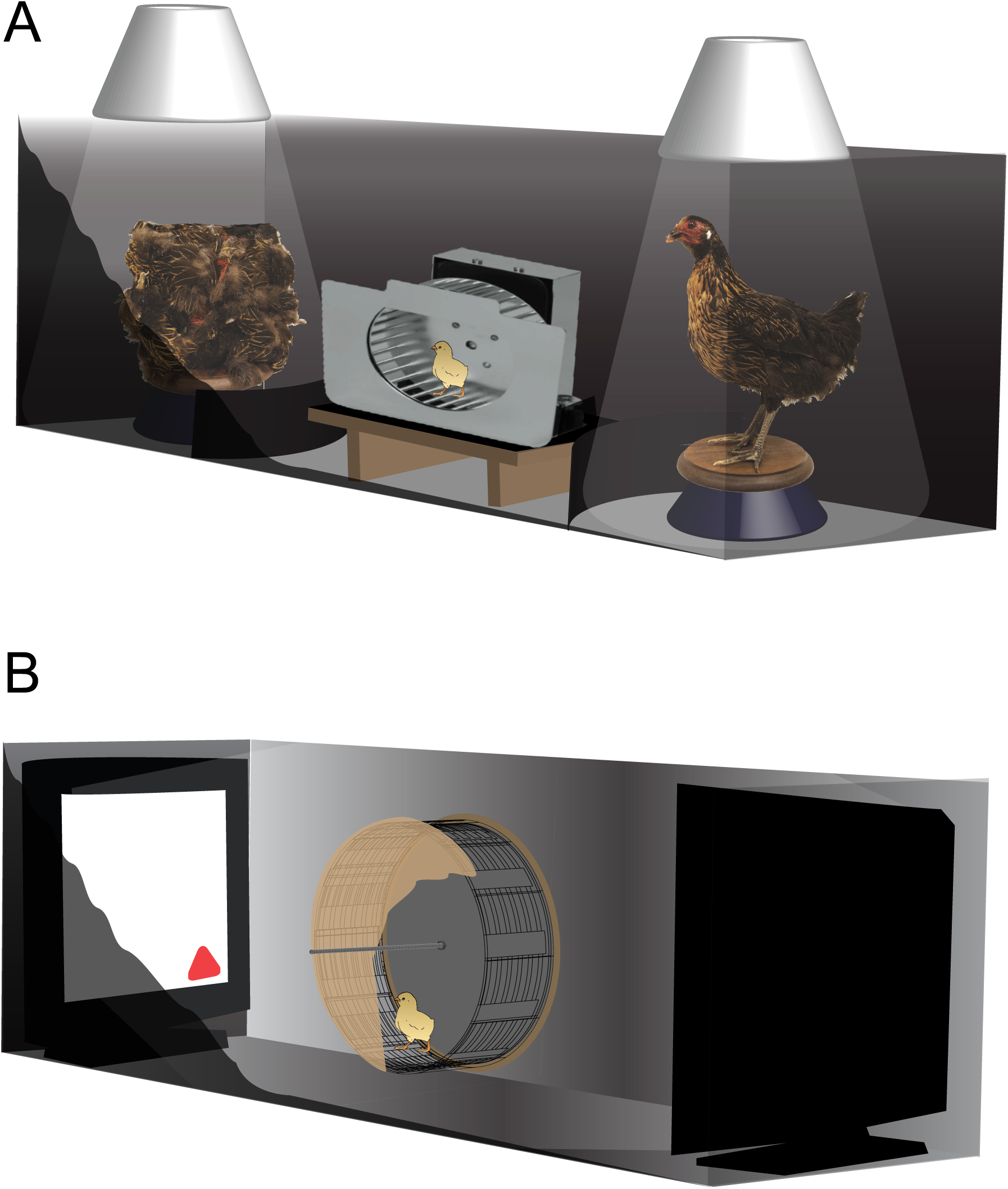
Experimental setup. Schematic representation of the (A) social predisposition and the (B) filial imprinting test apparatuses. In both cases the chick was placed in the running wheel and was free to approach either the social/familiar stimulus or the nonsocial/unfamiliar one, both visible at the two ends of the apparatus. The chick’s behavior was video-recorded from above. In (A) the stimuli consisted of a stuffed hen and a box, in which features of the hen were dismantled and attached on the sides of the box in a scrambled order [16,27,28,36,50], positioned on two rotating platforms and illuminated from above and by top/front lights (not shown here). In (B) imprinting stimuli were played on computer screens.

**Figure 2.**
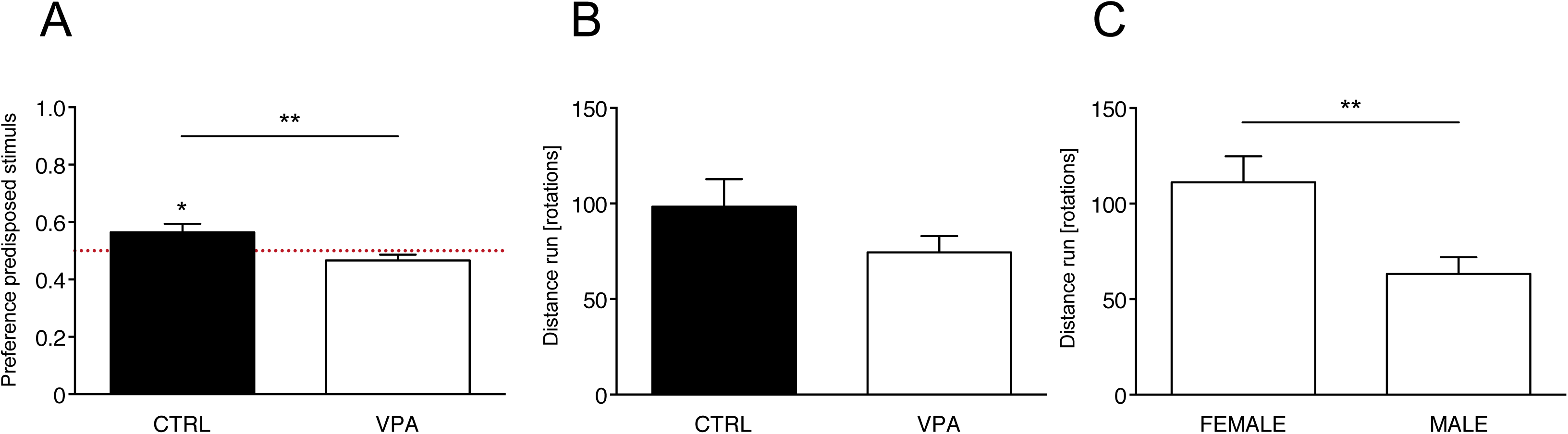
Social predispositions test. Bar graphs of preferences scores and motor activity in the social predispositions test. (A) Social preference test for stuffed hen (predisposed stimulus) and non-predisposed stimulus (see Methods for details). Analysis of variance of social preference scores using treatment and sex as between-subject factors, revealed a significant main effect of treatment (F_(1,58)_ = 7.708, p = .007; line with asterisks), with no other main effects or interactions among the other factors analyzed (sex (F_(1,58)_ = 0.026, p = .872), treatment x sex (F_(1,58)_ = .050, p = .823)). Preference scores were significantly different from chance level for the control group (CTRL: t_28_ = 2.191, p = .037), but not for VPA-treated chicks (VPA: t_32_ = −1.684, p = .102). Asterisks indicate significant departures from chance level, marked by the red line at 0.5. (B, C) Motor activity in the running wheel. Analysis of variance on number of rotations using treatment and sex as between-subject factors, showing (B) no significant main effects of treatment (F_(1,58)_ = 1.385, p = .244) or interaction treatment x sex (F_(1,58)_ = 0.021, p = .884) and (C) a significant effect of sex (F_(1,58)_ = 9.707, p = .003) independent of treatment. Data represent mean ± SEM, * p < 0.05; ** p < 0.01.

To understand whether VPA specifically affected social predispositions or impaired cognitive abilities and affiliative responses in general, we tested the effect of VPA also on filial imprinting. Differently from social predispositions, the learning mechanism of filial imprinting orients affiliative responses of chicks after previous exposure to a conspicuous stimulus [29,30]. Although it has been suggested that, in the wild, social predispositions might guide the learning process of filial imprinting by orienting the initial responses of chicks towards the mother hen, imprinting has a different neurobiological basis than social predispositions [31,32]. We exposed VPA- and vehicle-injected chicks to filial imprinting with artificial 2D objects (see Methods), and subsequently measured their learned preference for the imprinting stimulus *<u>vs. an</u>* unfamiliar stimulus. The chicks’ approach to the imprinting stimulus was not significantly different between treatment groups (Figure 3A), indicating no significant effect of VPA on the learning mechanisms of imprinting. Both groups successfully imprinted on both the stimuli, and no difference was detected in motor activity (Figure 3B).

**Figure 3.**
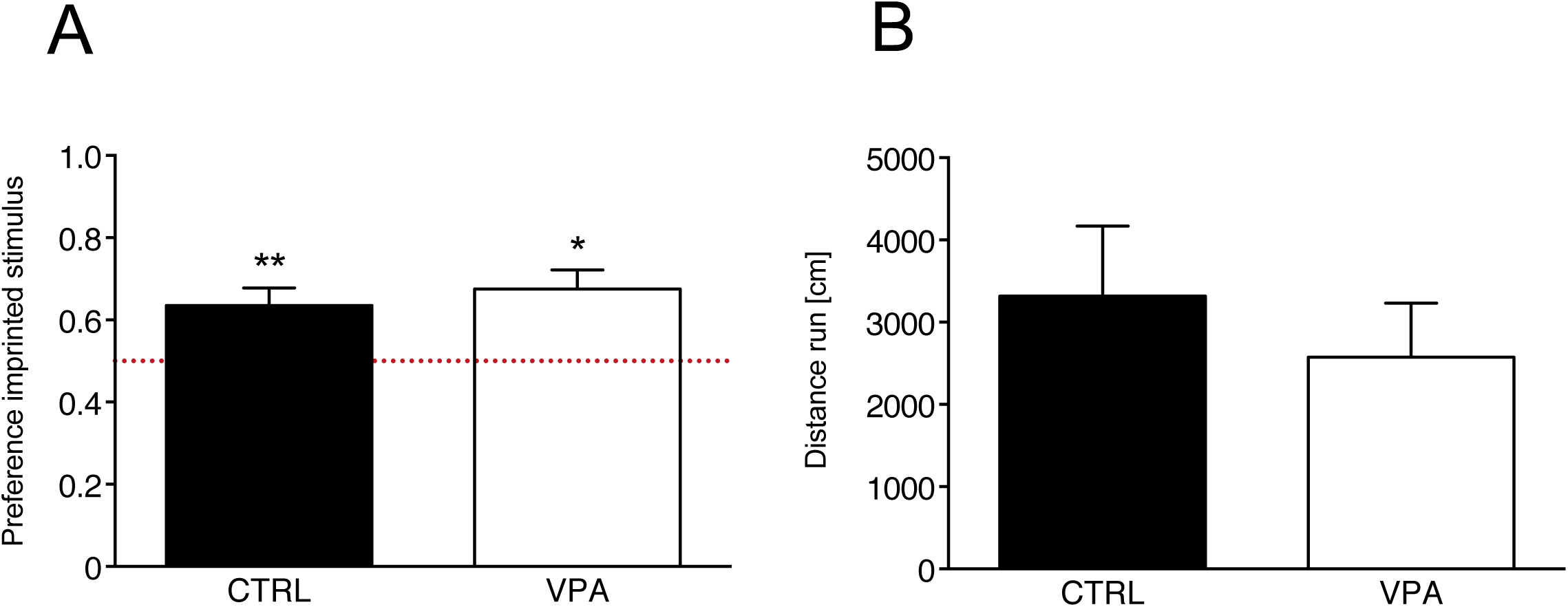
Imprinting test. Bar graphs of preferences scores and motor activity in the filial imprinting test. (A) Analysis of variance using treatment and sex as between-subject factors, revealed no significant main effect or interactions [treatment (F_(1,47)_ = .037, p = .849), sex (F_(1,47)_ = .009, p = .924) or treatment x sex (F_(1,47)_ = .331, p = .568)]. The mean preference scores for the imprinted stimulus were significantly different from chance level for both treatment groups (CTRL: t_25_ = 3.173, p = .004; VPA: t_28_ = 3.743, p = .001). Asterisks indicate significant departures from chance level, which is marked by the red line at 0.5. In both treatment groups preference scores > 0.5, indicating preference for the imprinted stimulus. (B) Analysis of variance on motor activity during the test did not detect any significant main effect of treatment (F_(1,47)_ = .666, p = .418), sex (F_(1,47)_ = .421, p = .520) or their interaction [treatment x sex (F_(1,47)_ = .000, p = .992)] in the motor activity of the chicks during the imprinting tests in the running wheel. Data represent mean ± SEM, * p < 0.05; ** p < 0.01.

## Discussion

Several evidence have demonstrated that early intervention can reduce the incidence and severity of symptoms in ASD children as young as 12 months of age [33-35]. To date, however, diagnosis of ASD is solely based on behavioral observations conducted after 2 years of age, limiting the possibility for early treatments. Given the importance of early diagnose and timely interventions for ASD patients, and the opportunity to shed light on the biological mechanisms underlying ASD and their developmental trajectories, the establishment of model that mimic the earliest behavioral alterations in ASD remains a focus of current research in the field. Precocial social species that exhibit early social predispositions at birth, such as domestic chicks, are a more convenient model to investigate these early-emerging social behaviors, compared to rodents. Using chicks to investigate ASD phenotypes, also allow to study the visual social predispositions observed in human neonates with striking similarities [2,6,7,9-12,18], to investigate the genetic bases for early predispositions [36], and to gain knowledge on the neurobiological bases of social predispositions [16,17,37] with an accurate control of the environment in which embryos and chicks develop [38]. In spite of this, to our knowledge, no work has been previously carried out on the possibility that VPA, a compound used to induce ASD-like behavioral deficits in many vertebrate species [39-43], may affect early social predispositions. Here we combined the evolutionary conserved effect of VPA on brain development [39-42] with the analysis of specific social behaviors common to human newborns and newly-hatched chicks.

Our results show a detrimental effect of VPA on the well documented predispositions to approach a social stimulus (stuffed hen) over a comparable “non-social” stimulus (scrambled hen-like box) [27,28]. Previous studies showed that chicks’ preferences are elicited by the presence of the head and neck region [44]. Indeed, faces and face-like configuration cues, analogous to those present in the stuffed hen stimulus, are known to elicit a strong preference in both chicks and human neonates [4,6].

On the contrary, when we investigated affiliative responses mediated by the experience-dependent mechanism of filial imprinting, we found that VPA did not impair these responses. These results indicate that, despite the fact that VPA disrupted the chicks’ predisposition to preferentially approach the stuffed hen, the brain capacity to activate experience-dependent learning mechanisms remains intact. This data reveals a specific effect of VPA on social behaviors [26] and in particular, on social predispositions. Indeed, despite its wide biological targets, acute exposure to VPA during late embryogenesis has been shown to act preferentially on social behaviors, and to spare general cognitive and motor functions [45].

Accumulating evidence suggest that the deficits in social behavior underlying autism spectrum disorder (ASD) may be the result of poor orienting and attention to important social stimuli, such as faces and face-like configuration cues, during early infancy [46-48]. Since social predispositions represent the earliest emerging social behaviors in humans, impairments in this domain could result in limited interests for relevant social stimuli at the time of birth [49], preventing neonates from focusing their attention on salient social stimuli, compromising the typical developmental trajectories of the social brain and contributing to the appearance of ASD symptoms [4]. Recent data showed impairments in these orienting mechanisms in human neonates at high-risk for ASD [18]. Our data further confirm this hypothesis in an animal model, showing the suppression of a predisposed social preference in domestic chicks exposed to VPA. Furthermore our data may indicate that VPA treatment in domestic chicks affects development of the social brain circuits at the base of social predisposition, which could be involved in the earliest expression of ASD symptoms. In fact, recent studies demonstrated a response of some nodes of the “social behavior” and “social decision-making” networks during the very first exposure of visually naïve chicks to conspecifics, or to some elementary visual properties that elicit their spontaneous social preferences [17]. This data suggests that development and tuning of those networks crucial for the control of adult social behavior, might be shaped by early visual inputs that the organisms receives from social companions, thanks to its social predispositions and to the species-typical structure of the environment.

Altogether, we believe that our study has a very high translational value; opening new perspectives to investigate the mechanistic bases of ASD with the aid of behavioral markers analogous to those used in human neonates studies. Further studies should clarify whether VPA effect extends on other social predispositions, such as preferences for dynamic stimuli that change in speed [13,28], whether this substance has a disruptive or a delaying effect on social predispositions [50] and on possibilities to rescue social responses after exposure to VPA.

## Methods

### Ethics statement

All experiments comply with the current Italian and European Community laws for the ethical treatment of animals, and the experimental procedures were approved by the Ethical Committee of the University of Trento and licensed by the Italian Health Ministry (permit number 986/2016-PR).

### Chick embryo injections

Freshly fertilized eggs of domestic chicks (*Gallus gallus*), of the Ross 308 (Aviagen) strain, were obtained from a local commercial hatchery (Agricola Berica, Montegalda (VI), Italy), placed in a cold room at 4 °C and maintained in a vertical position for 24-72 h. The eggs were then placed in the dark and incubated at 37.5 °C and 40% relative humidity, with rocking. The first day of incubation was considered embryonic day 0 (E0). Fertilized eggs were then selected by a light test on E14 and injected. Chick embryo injection was performed according to previous reports [26]. Briefly, a small hole was made on the egg shell above the air sac, and 35 µmoles of VPA (Sodium Valproate, Sigma Aldrich) were administered, in a volume of 200 µl, to each fertilized egg, by dropping the solution onto the chorioallantoic membrane. Age-matched control eggs were injected using the same procedure with 200 µL of vehicle (double distilled injectable water). After sealing the hole with paper tape, eggs were placed in a rocking incubator (FIEM srl, Italy) until E18, when eggs were placed in a hatching incubator (FIEM srl, Italy). Hatching took place at a temperature of 37.7 °C, with 60% humidity, as previously described [16]. The day of hatching was considered post-hatching day 0 (P0). All subsequent procedures were performed in complete darkness, so that the chicks remained visually inexperienced until the moment of test.

## Experiment 1. Social Predisposition test

### Rearing conditions, apparatus and stimuli

We used the same procedure previously described to assess chicks’ social predispositions [51]. At P1 (24 h after hatching), chicks were transferred to individual compartments (11 cm × 11 cm × 25 cm) at the constant temperature of 33 °C. To enhance the expression of the social predispositions, chicks were exposed to acoustic stimulation [51] inside a dark incubator equipped with a loudspeaker. Non-species-specific sound stimulation was provided using a digitally constructed audio file as previously described [16]. The test apparatus consisted of a running wheel mounted at the center of a 150cm-long and 46cm-wide arena, with lateral walls of 45 cm of height [27]. Stimuli were located at the opposite sides of the apparatus, on two rotating platforms (30 rotations per minute), illuminated from above (40 W warm diffused light) and by top/front lights (25 W warm light). The test stimuli consisted in an intact jungle fowl hen and box, in which features of the hen were dismantled and attached on the sides of the box in a scrambled order, as described previously [16,27,28,36,52].

### Test procedure

Chicks’ preferences for a stuffed hen *vs.* a scrambled stuffed hen were tested at P2 for a 30 minutes. Each subject was individually extracted from the incubator in complete darkness and carried, in a closed box, to the experimental room. At the beginning of the test chicks were individually placed in the running wheel facing one of the lateral walls, so that they could see both stimuli on the opposite sides of the apparatus. Their approach responses were recorded as distance run (number of wheel rotations) in the direction of each stimulus. Right/left stimulus position, sex and treatment were balanced between experimental sessions. Each session was video recorded.

## Experiment 2. Filial imprinting test

### Rearing conditions, apparatus and stimuli

Chicks hatched in individual compartments (11 x 8.5 x 14 cm) in complete darkness. Chicks were placed in the imprinting set-up soon after hatching, with water and food *ad libitum*, and exposed to the imprinting stimulus for 3 days. The imprinting set-up consisted of a black box (30 cm x 30 cm x 40 cm) with a monitor (17”, 60 Hz) mounted on the front wall, displaying one imprinting visual stimulus for 14 hours a day, continuously. During the remaining 10 hours the screen was black and the lights were off, allowing the normal day-night cycle. Chicks were exposed to either a blue circle or a red triangle (3.7 cm circle diameter and triangle sides), moving at 1.5 cm/s on a white screen [53,54]. The same stimuli (the imprinting stimulus and the unfamiliar stimulus, that the chicks had never seen before) were used for imprinting test. Chicks were individually tested at P3. The imprinting test apparatus consisted of a 150 cm long, 46 cm wide, 45 cm high arena, equipped with a running wheel (32 cm diameter, 13 cm large, covered with 1 cm of opaque foam on both sides) in the center of the apparatus.

### Test procedure

Chicks’ preferences for the imprinting stimulus (familiar) *vs*. an unfamiliar stimulus were tested at P3 for 20 minutes. Chicks were individually placed in the center of the running wheel, facing one of the lateral walls, so that they could see both stimuli on the opposite sides of the apparatus. The preference for the imprinting and for the unfamiliar stimulus was measured using the distance run (centimeters) towards each stimulus. Right/left stimulus position, imprinting stimulus (red triangle or blue circle), sex and treatment were balanced between experimental sessions. Each session was video recorded.

### Statistical analysis

To assess social predispositions and imprinting responses independently from motor activity, we calculated for each chick a preference score for the predisposed/imprinted stimulus adjusted for the overall distance run, as

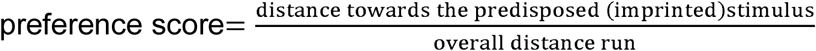

Values of this ratio range from 1 (full choice for the social stimulus) to 0 (full choice for the non-social stimulus), where 0.5 represents the absence of preference. We assessed differences in the motor activity by comparing the overall distance run, regardless of the approached stimulus, for the entire test session. Effect of treatment, sex and type of imprinted stimulus on the preference score was evaluated by multifactorial analysis of variance (ANOVA). For post-hoc analysis, *t*-tests were used. For all the tests, significant departures of the preference score from chance level (0.5) were estimated by one-sample two-tailed *t*-tests. All statistical analyses were performed with IBM SPSS Statistic for Windows (Version 24.0). Alpha was set to 0.05 for all tests.

## Acknowledgments

This work was supported by a grant from the European Research Council under the European Union’s Seventh Framework Programme (FP7/2007-2013)/Advanced Grant ERC PREMESOR G.A. [n 295517] to GV. Support from Fondazione Caritro Grant Biomarker DSA [40102839] is also acknowledged.

